# Draft genomes of a male and female Australian jacky dragon (*Amphibolurus muricatus*)

**DOI:** 10.1101/2021.10.11.463868

**Authors:** Ran Tian, Hao Dong, Fan Zhang, Hao Yu, Enqing Pei, Chengcheng Shi, Guangyi Fan, Sarah L. Whiteley, Clare E. Holleley, Inge Seim, Arthur Georges

**Author notes:** Corresponding author: Institute for Applied Ecology, University of Canberra, ACT 2601, Australia. These authors contributed equally to this work and share first authorship (Ran Tian, Hao Dong).

## Abstract

Australia is remarkable for its lizard diversity, with very high endemicity because of continental-scale diversification and adaptive radiation during prolonged isolation. We here employed stLFR linked-read technology to generate male and female draft genomes of the jacky dragon *Amphibolurus muricatus*, an Australian dragon lizard (family Agamidae; the agamids). The assemblies are 1.8 Gb in size and have a repeat content (39%) and GC content (42%) similar to other dragon lizards. The longest scaffold was 39.7 Mb (female) and 9.6 Mb (male), with corresponding scaffold N50 values of 6.8 Mb and 1.6 Mb. The BUSCO (Sauropsida database) completeness percentages were 90.2% and 88.8% respectively. Phylogenetic comparisons show that Australian and Asian agamids split from a common ancestor about 80 million years ago, while the Australian genera *Amphibolurus, Pogona*, and the basal *Intellagama* split ∼37 million years ago. The draft *A. muricatus* assemblies will be a valuable resource for understanding lizard sex determination and the evolution and conservation of Australian dragon lizards.

## Introduction

The Australian jacky dragon *Amphibolurus muricatus* (**Figure 1**) is a lizard that is widespread in dry sclerophyll forests of south-eastern and eastern Australia (Cogger 2014). It is a model species for biogeography (Pepper et al. 2014), evolutionary biology (Warner et al. 2013; Warner and Shine 2008), social behaviour (Peters and Evans 2003; Woo and Rieucau 2013) and development (Whiteley et al. 2021; Esquerré et al. 2014).

**Figure 1.**
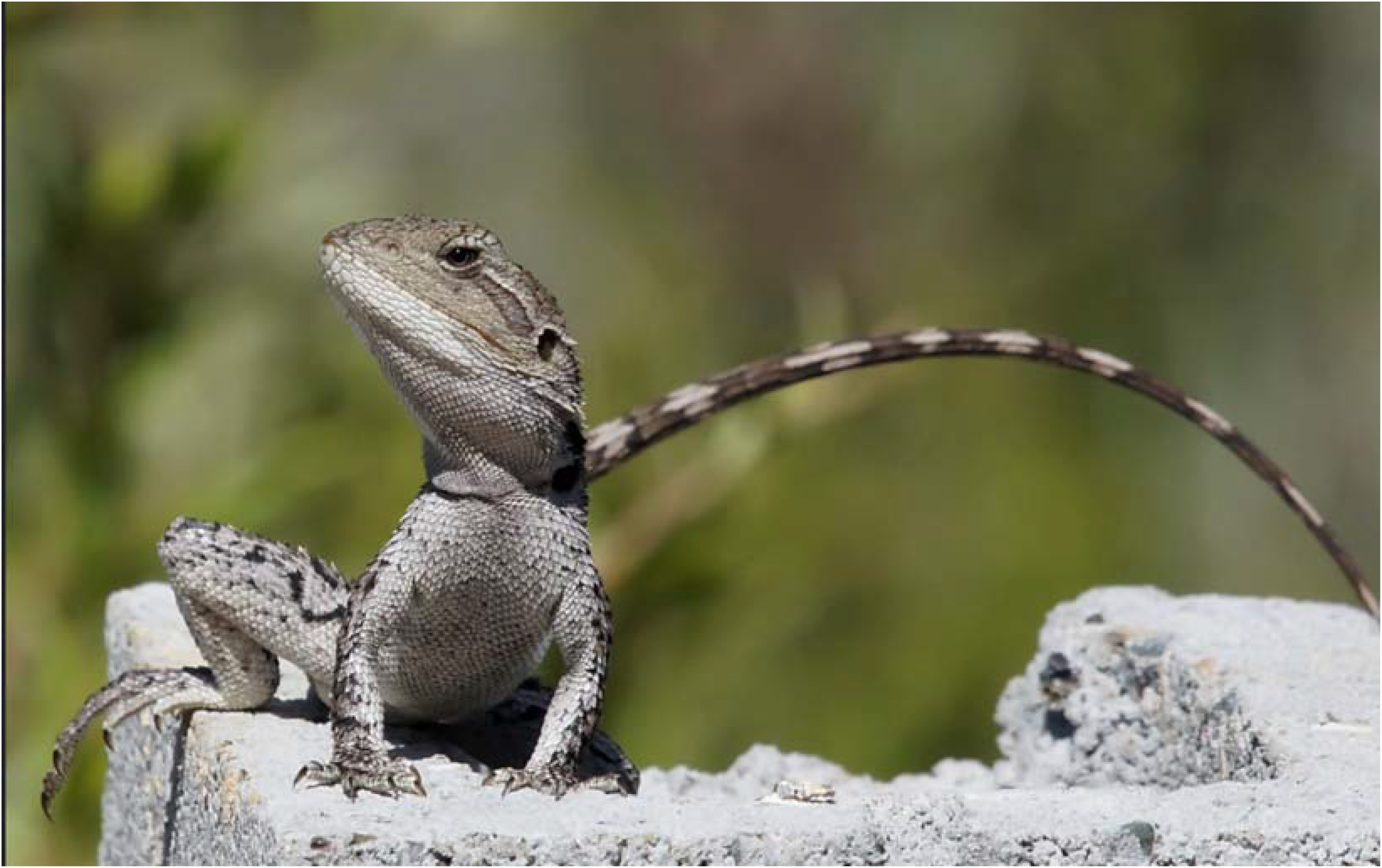
Photograph of an adult male jacky dragon (*Amphibolurus muricatus*). Image credit: David Cook Wildlife Photography.

Species in the genus *Amphibolurus* and *Chlamydosaurus* are a major clade in the Australian radiation of the Agamidae (Hugall et al. 2008). The draft assembly of *A. muricatus*, together with that of *Pogona vitticeps* (Georges et al. 2015), represents the first foray into generating the necessary high-quality genomes for the Agamidae. In particular, *A*.*muricatus* occupies mesic habitats and so is intermediate between the Australian water dragon *Intellagama lesueurii* and the forest dragon *Lophosaurus boydii* that occupy hydric habitats, and the central bearded dragon *Pogona vitticeps* and the Lake Eyre dragon *Ctenophorus maculosus*, for example, that occupy more xeric habitats. As such, it is one of several species important for understanding genomic adaptation to the progressive aridity that has occurred in Australia in the past 15 Myr. *Amphibolurus muricatus* is also of particular interest because it has temperature-dependent sex determination (TSD) (Harlow and Taylor 2000) and it is unclear as to whether this arises from classical TSD or a combination of genetic and environmental influences (Whiteley et al. 2021). Studies of the underlying mechanisms of TSD require a genome assembly and knowledge of genome organisation to identify genes on the sex chromosomes of species with genotypic sex determination (GSD) and their chromosomal and gene homology in closely related TSD species. This is particularly so in species with TSD that show evidence of cryptic residual or de novo genotypic influence on offspring sex ratios, as is suspected for *A. muricatus* (Whiteley et al. 2021).

Here, we generated draft, annotated genome assemblies for a male and a female *A. muricatus*. We used transcriptomes sequenced and assembled for *A. muricatus* and published genomes (*Anolis, Varanus* and *Pogona*) to annotate the assemblies. Our assemblies will provide a resource to increase capacity and accelerate the progress of studies into the evolution, ecology, and conservation of Australian dragon lizards.

## Materials and methods

### Sample collection

In attempt to reduce the high heterozygosity that presented difficulties in the assembly of the genome of *Pogona vitticeps* (Georges et al., 2015), we generated inbred lines of *A. muricatus*. The founding male and female pair were sourced from the wild and bred in captivity. The two animals used to generate the genome were obtained from the fourth generation of the inbred pedigree produced by sib-sib matings and back crossing (see **Figure S1** for the complete pedigree). The male (AA069033) and female (AA069032) individuals used for the genome and transcriptome sequencing were humanely euthanised via intraperitoneal injection of sodium pentobarbitone (60 mg/ml in isotonic saline). Organs were rapidly dissected and snap frozen in liquid nitrogen.

### DNA extraction

High molecular weight DNA was extracted from liver (female) and blood (male). DNA for 10x Genomics linked-read sequencing was extracted using a QIAGEN Gentra Puregene DNA Isolation kit (to extract >100 kb DNA fragments). DNA for stLFR linked-read sequencing was extracted using a QIAGEN MagAttract HMW DNA Kit (to extract >40 kb DNA fragments). DNA yield and quality was assessed using a NanoDrop spectrophotometer (Thermo Fisher Scientific, Waltham, MA, USA) and a Qubit fluorometer (Thermo Fisher Scientific) and pulse-field gel electrophoresis.

### Genome size and heterozygosity estimation

We used the paired-end stLFR sequencing reads to estimate the genome size and heterozygosity of the female and male *A. muricatus* samples. Briefly, *k*-mers (*k*=17, as employed in a previous study on *P. vitticeps* (Georges et al. 2015)) in reads were counted and converted to histogram files using Jellyfish (v2.2.10) (Marcais and Kingsford 2011), followed by analysis using GenomeScope v2.0 (Vurture et al. 2017; Ranallo-Benavidez et al. 2020).

### Assembly 1.0: A 10x Genomics linked-read sequencing assembly

Male and female *A. muricatus* genome sequencing libraries were constructed on the Chromium system (10x Genomics, Pleasanton, CA, USA) by the Ramaciotti Centre for Genomics (Sydney, Australia). The Chromium instrument enables unique barcoding of long stretches of DNA on gel beads. The barcodes allow later reconstruction of long DNA fragments from a series of short DNA fragments with the same barcode (i.e., linked-reads). After barcoding, DNA was sheared into smaller fragments and sequenced on the NovaSeq 6000 platform (Illumina, CA, USA) to generate 151 bp paired-end (PE) reads. A total of 904.9 M raw 10x Genomics Chromium linked-reads were generated. Raw 10x data were assembled with Supernova v2.1.1 (Weisenfeld et al. 2017) and a FASTA file was generated using the ‘pseudohap style’ option in Supernova mkoutput. All female (∼450 M) and male (∼550 M) read pairs were used (female sequencing depth *ca* 50.3×; male, *ca* 47.8×). The resulting assemblies were further scaffolded with ARKS v1.0.3 (Coombe et al. 2018), reusing the 10x reads and the companion LINKS program (v1.8.7) (Warren et al. 2015). ARKS employs a *k*-mer approach to map linked barcodes to the contigs in the initial Supernova assembly to generate a scaffold graph with estimated distances for LINKS input. These assemblies were denoted AmpMurF_1.0 (female) and AmpMurM_1.0 (male). We used GapCloser v1.12 (part of SOAPdenovo2) (Luo et al. 2012) to fill gaps in the assembly. GapCloser was run using the parameter -l 150) and clean 10x Genomics reads PE reads.

### Assembly 1.1: Further scaffolding of assembly 1.0 using RNA-seq data

We attempted to improve the v1.0 genome assemblies’ contiguity using RNA-sequencing reads. RNA-seq reads (from brain, ovary, and testis; see below) were filtered (i.e., cleaned) to remove adapters and low-quality reads using Flexbar v3.4.0 and used to further re-scaffold the v1.0 assemblies (FASTA files before gapclosing) with P_RNA_scaffolder (Zhu et al. 2018). The default Flexbar settings discards all reads with any uncalled bases. A final round of scaffolding was performed on the resulting assemblies using L_RNA_scaffolder (Xue et al. 2013). These assemblies were denoted AmpMurF_1.1 (female) and AmpMurM_1.1 (male). As before, GapCloser and clean 10x Genomics reads were used to fill gaps.

### Assembly 2.0: Further scaffolding of assembly 1.0 using SLR-superscaffolder

As an alternative approach, we attempted to improve the v1.0 genome assemblies’ contiguity using SLR-superscaffolder (Guo et al. 2021). Briefly, SLR-superscaffolder employs single-tube long fragment read (stLFR) sequencing (Wang et al. 2019) reads (see section below) to generate hybrid genome assemblies. The software was run with default parameters except for PE_SEED_MIN=300 (minimum contig size to fill; default 1000). These assemblies were denoted AmpMurF_2.0 (female) and AmpMurM_2.0 (male). GapCloser and clean stLFR reads (with the barcode removed using https://github.com/BGI-Qingdao/stLFR_barcode_split) were used to fill gaps.

### Assembly 3.0: An stLFR linked-read sequencing assembly

We also generated independent assemblies for the individuals sequenced on the 10x Genomics Chromium system using stLFR sequencing (Wang et al., 2019). BGI (Brisbane, Australia) generated ∼100× coverage 100-bp paired-end reads (plus a 42-bp stLFR barcode on the right/_2 read) per individual. Low-quality reads, PCR duplicates, and adaptors were removed using SOAPnuke v1.5 (Chen et al. 2018). All female (∼1,517 M) and male (∼1,427 M) read pairs were utilised. The stLFRdenovo pipeline (https://github.com/BGI-biotools/stLFRdenovo), which is based on Supernova v2.11 and customised for stLFR data, was used to generate a *de novo* genome assembly. The stLFRdenovo tool ‘FillGaps’ was used to fill gaps.

### RNA-seq and transcriptome assembly

Raw data 125 bp PE reads, generated on an Illumina HiSeq 2500 instrument was filtered using Flexbar v3.4.0 (Roehr et al. 2017; Dodt et al. 2012) with default settings (eliminates reads with any uncalled bases). Any residual ribosomal RNA reads (the majority removed by poly(A) selection prior to sequencing library generation) were removed using SortMeRNA v2.1b (Kopylova et al. 2012) against the SILVA v119 ribosomal database (Quast et al. 2013). Tissue transcriptomes were de novo assembled using Trinity v2.11.0 (Haas et al. 2013; Grabherr et al. 2011; Henschel et al. 2012) and assessed using BUSCO v5.0.0.

### Genome annotation

We identified repetitive elements by integrating homology and de novo prediction data. Protein-coding genes were annotated using homology-based prediction, de novo prediction, and RNA-seq-assisted prediction methods.

Homology-based transposable elements (TE) annotations were obtained by interrogating a genome assembly with known repeats in the Repbase database v16.02 (Bao et al. 2015) using RepeatMasker v4.0.5 (DNA-level) (parameters: -species lizard -e rmblast - xsmall -s -gff -pa 4) (Tarailo-Graovac and Chen 2009) and RepeatProteinMask (protein-level; implemented in RepeatMasker). De novo TE predictions were obtained using RepeatModeler v1.1.0.4 (parameters: -pa 4 -database lizard -LTRStruct) (Smit and Hubley 2010) and LTRharvest v1.5.8 (default parameters) (Ellinghaus et al. 2008) to generate database for a RepeatMasker run. Tandem Repeat Finder v4.07 (2 7 7 80 10 50 500 -m -f -d) with default settings (i.e., MaxPeriod parameter, the maximum period size,) (Benson 1999) was used to find tandem repeats (TRs) in the genome. A non-redundant repeat annotation set was obtained by combining the above data.

Protein-coding genes were annotated using homology-based prediction, de novo prediction, and RNA-seq-assisted [generated from ovary, testis, and brain (both sexes)] prediction methods. Sequences of homologous proteins from three lizards [*Anolis carolinensis* (green anole) assembly AnoCar2.0 (RefSeq assembly GCF_000090745.1) (Alfoldi et al. 2011); *Varanus komodoensis* (Komodo dragon) assembly ASM479886v1 (GCA_004798865.1) (Lind et al. 2019); and *Pogona vitticeps* (central bearded dragon) assembly pvi1.1 (GCF_900067755.1)] (Georges et al. 2015) were downloaded from NCBI. These protein sequences were aligned to the repeat-masked genome using BLAT v0.36 (Kent 2002). GeneWise v2.4.1 (Birney et al. 2004) was employed to generate gene structures based on the alignments of proteins to a genome assembly. De novo gene prediction was performed using AUGUSTUS v3.2.3 (Stanke et al. 2006), GENSCAN v1.0 (Burge and Karlin 1997), and GlimmerHMM v3.0.1 (Majoros et al. 2004) with a human training set. Transcriptome data (clean reads) were mapped to the assembled genome using HISAT2 v2.1.0 (Kim et al. 2019) and SAMtools v1.9 (Li et al. 2009), and coding regions were predicted using TransDecoder v5.5.0 (Grabherr et al. 2011; Haas et al. 2013). A final non-redundant reference gene set was generated by merging the three annotated gene sets using EvidenceModeler v1.1.1 (EVM) (Haas et al. 2008) and excluding EVM gene models with only ab initio support. The gene models were translated into amino acid sequences and used in local BLASTp (Camacho et al. 2009) searches against the public databases Kyoto Encyclopedia of Genes and Genomes (KEGG; v89.1) (Kanehisa and Goto 2000), NCBI non-redundant protein sequences (NR; v20170924) (O’Leary et al. 2016), Swiss-Prot (release-2018_07) (UniProt Consortium 2012), and InterPro (v69.0) (Mitchell et al. 2019).

### Phylogeny and divergence time estimation

In addition to *A. carolinensis, V. komodoensis* and *P. vitticeps* (see section above), the genome and sequences of homologous proteins from *Gekko japonicus* (Schlegel’s Japanese gecko) assembly Gekko_japonicus_V1.1 (GCA_001447785.1) (Liu et al. 2015) and *Crotalus tigris* (tiger rattlesnake) assembly ASM1654583v1 (GCA_016545835.1) (Margres et al. 2021) were downloaded from NCBI. The genome and annotations of *Ophisaurus gracilis* (Anguidae lizard) were downloaded from GigaDB (Song et al. 2015a; Song et al. 2015b). No gene annotation data were available for three species: *Intellagama lesueurii* (Australian water dragon; assembly EWD_hifiasm_HiC generated as part of the AusARG consortium and (downloaded from DNA Zoo (Dudchenko et al. 2018; Cheng et al. 2021; Dudchenko et al. 2017)) and the Chinese agamid lizards *Phrynocephalus przewalskii* (Przewalski’s toadhead agama) (Gao et al. 2019) and *Pyrocephalus vlangalii* (Ching Hai toadhead agama) (Gao et al. 2019) (CNGBdb accession no. CNP0000203). Their protein-coding genes were annotated using homology-based prediction, de novo prediction, and RNA-seq-assisted prediction methods (see genome annotation section above).

We identified 1:1 orthologs by interrogating the predicted proteins from the gene models of 13 squamate species using SonicParanoid v1.3.0 (Cosentino and Iwasaki 2019). The corresponding coding sequences (CDS) for each species were aligned using PRANK v100802 (Loytynoja and Goldman 2005) and filtered by Gblocks v0.91b (Talavera and Castresana 2007) to identify conserved blocks (removing gaps, ambiguous sites, and excluding alignments less than 300 bp in size), leaving 3,130 genes. Maximum-likelihood (ML) phylogenetic trees were generated using RaxML v7.2.8 (Stamatakis 2006) and IQ-Tree v2.1.3 (Minh et al. 2020) with three CDS data sets: the whole coding sequence (whole-CDS), first codon positions, and fourfold degenerate (4d) sites. Identical topologies and similar support values were obtained (1,000 bootstrap iterations were performed). The divergence time between species was estimated using MCMCTree [a Bayesian molecular clock model implemented in PAML v4.7 (Yang 2007)] with the JC69 nucleotide substitution model, and the whole-CDS ML tree and concatenated whole-CDS supergenes as inputs. We used 100,000 iterations after a burn-in of 10,000 iterations. MCMCTree calibration points (million years ago; Mya) were obtained from (Oliver and Hugall 2017) (crown age of Australian agamids 27.1 Mya, with 95% CI 20.1-37.7) and TimeTree (Kumar et al. 2017): *G. japonicus-P. przewalskii* (190–206 Mya), *V. komodoensis-O. gracilis* (121–143 Mya), *V. komodoensis-C. tigris* (156–174 Mya), *V. komodoensis-A. carolinensis* (155–175 Mya), *I. lesueurii-A. carolinensis* (139–166 Mya), *I. lesueurii-P. przewalskii* (73–93 Mya), *I. lesueurii-A. muricatus* (25.5–42.4 Mya), *P. vitticeps-A. muricatus* (20.2–34.6 Mya).

## Results and discussion

### Draft genome assembly and comparisons with other squamates

We used stLFR reads, Jellyfish (Marcais and Kingsford 2011), and GenomeScope (Vurture et al. 2017; Ranallo-Benavidez et al. 2020) to estimate genome size and heterozygosity. The size of the *A. muricatus* genome was estimated to be around 1.5 Gb. The genome-wide heterozygosity from our inbred (four generations) *A. muricatus* lines was estimated to range from 0.66% (female) to 0.73% (male), slightly lower than the central bearded dragon (*Pogona vitticeps*) (0.85%) (Georges et al. 2015).

We generated four genome assemblies per sample. The v1.0 assemblies were generated using 10x Genomics Chromium data and the Supernova assembler and further refined using ARKS and LINKS. The v1.1 assemblies employed P_RNA_scaffolder (uses RNA-seq reads from brain, ovary, and testis) (**Table S1**) and L_RNA_scaffolder (uses Trinity transcriptome assemblies) (**Tables S2** and **S3**) to improve the v1.0 assemblies, while the v2.0 assemblies used SLR-superscaffolder and stLFR reads to improve the v1.0 assemblies. Finally, the 3.0 assemblies were generated using stLFR reads alone and Supernova. While re-scaffolding of the assemblies generated using 10x Genomics Chromium sequencing improved the initial v1.0 assembly (in particular, SLR-superscaffolder), assembly using stLFR data alone gave the best assembly result (**Table S4**).

For the remainder of the manuscript, conclusions will be drawn from the *A. muricatus* v3.0 assemblies. A comparison with seven other agamid genome assemblies is shown in **Table 1**. The final, v3.0 assemblies have a total scaffold length (i.e., containing gaps) of 1.87 (female) and 1.80 (male) Gb. The longest scaffold was 39.7 Mb (female; AmpMurF_3.0) and 9.6 Mb (male; AmpMurM_3.0), and the corresponding scaffold N50 values of 6.9 and 1.6 Mb. The contig N50s were 67.2 kb (AmpMurF_3.0) and 59.3 kb (AmpMurM_3.0). The N50 values are lower than most recent squamate assemblies but similar to the agamids central bearded dragon (*P. vitticeps*; scaffold N50 2.29 Mb and contig N50 31.30 kb) (Georges et al. 2015), Przewalski’s toadhead agama (*P. przewalskii*; scaffold N50 6.88 MB and contig N50 56.43 kb) (Gao et al. 2019), and Ching Hai toadhead agama (*P. forsythii*; scaffold N50 2.40 Mb and contig N50 31.29 kb) (Gao et al. 2019) (**Figure 2**). The BUSCO (v5.0.0 with the 7,480-gene sauropsida_odb10 dataset) metrics of the *A. muricatus* assemblies compared well to other squamate assemblies, including agamids from Australia [*P. vitticeps* (Georges et al. 2015) and *I. lesueurii* (Australian water dragon)], China (toad-headed agamas of genus *Phrynocephalus* sp. (Gao et al. 2019; Jin et al. 2022; Qi et al. 2023)), and the widely distributed oriental garden lizard (*C. versicolor*) (Wang et al. 2023) (**Figure 3** and **Table S5**).

**Table 1.** Agamidae genome assembly statistics. Lengths in base pairs (bp). AmpMurF_3.0 and AmpMurM_3.0 denotes the female and male *Amphibolurus muricatus* assemblies, respectively. NA denotes not applicable. Assemblies with the suffix _HiC are available at DNA Zoo; CalVer_1.0 (CNP0003598), Phprz_v1.0 (CNP0000203), and Phvla_v1.0 (CNP0000203) are available at the China National GeneBank Database; the remainder are available via NCBI.

| Species name | *Amphibolurus muricatus* | *Amphibolurus muricatus* | *Pogona vitticeps* | *Intellagama lesueurii* | *Phrynocephalus przewalskii* | *Phrynocephalus vlangalii* | *Phrynocephalus forsythii* | *Phrynocephalus versicolor* | *Calotes versicolor* |
| --- | --- | --- | --- | --- | --- | --- | --- | --- | --- |
| Common name | Jacky dragon | Jacky dragon | Central bearded dragon | Australian water dragon | Przewalski's toadhead agama | Ching Hai toadhead agama | Moustache toadhead agama | Variegated toadhead agama | Oriental garden lizard |
| Assembly | AmpMurF_3.0 | AmpMurM_3.0 | pvi1.1 | EWD_hifiasm_HiC | Phprz_v1.0 | Phvla_v1.0 | ASM2928247v1 | PhrVer | CalVer_1.0 |
| Reference | This study | This study | Georges *et. al* (2015) | NA | Gao *et al.*, 2019 | Gao *et al.*, 2019 | Qi *et al.* 2023 | Jin *et al.* 2022 | Wang *et al.* 2023 |
| Contig number | 145,095 | 95,472 | 636,780 | 1,457 | 592,043 | 506,434 | 944 | 102,689 | 104 |
| Contig length | 1,804,035,661 | 1,752,355,218 | 1,747,541,145 | 1,816,116,528 | 1,738,531,458 | 1,773,991,985 | 1,816,858,791 | 1,592,519,988 | 1,614,101,240 |
| Contig N50 (bp) | 67,166 | 59,294 | 31,297 | 11,213,063 | 56,431 | 31,286 | 46,215,561 | 27,024 | 91,599,063 |
| Contig max length | 645,479 | 773,372 | 295,776 | 55,971,000 | 547,676 | 368,251 | 95,027,530 | 275,221 | 160,793,834 |
| Scaffold number | 97,556 | 45,762 | 545,310 | 619 | 543,179 | 411,697 | 644 | 4,557 | 104 |
| Scaffold length | 1,868,109,324 | 1,804,786,947 | 1,816,116,151 | 1,816,535,528 | 1,763,820,185 | 1,820,170,410 | 1,816,888,593 | 1,603,387,288 | 1,614,101,240 |
| Scaffold N50 (bp) | 6,818,063 | 1,568,728 | 2,290,546 | 268,945,440 | 6,884,165 | 2,396,942 | 133,255,571 | 49,226,030 | 91,599,063 |
| Scaffold max length | 39,679,044 | 9,582,854 | 14,681,335 | 353,375,234 | 37,740,249 | 13,874,271 | 212,878,235 | 102,350,298 | 160,793,834 |
| Gaps (bp) | 64,073,663 | 52,431,729 | 68,575,006 | 419,000 | 25,288,727 | 46,178,425 | 29,802 | 10,867,300 | 0 |
| Gaps (%) | 3.43 | 2.91 | 3.78 | 0.02 | 1.43 | 2.54 | 0.0016 | 0.68 | - |
| GC content (%) | 41.75 | 41.67 | 41.87 | 41.94 | 42.94 | 43.07 | 43.32 | 42.91 | 43.79 |

**Figure 2.**
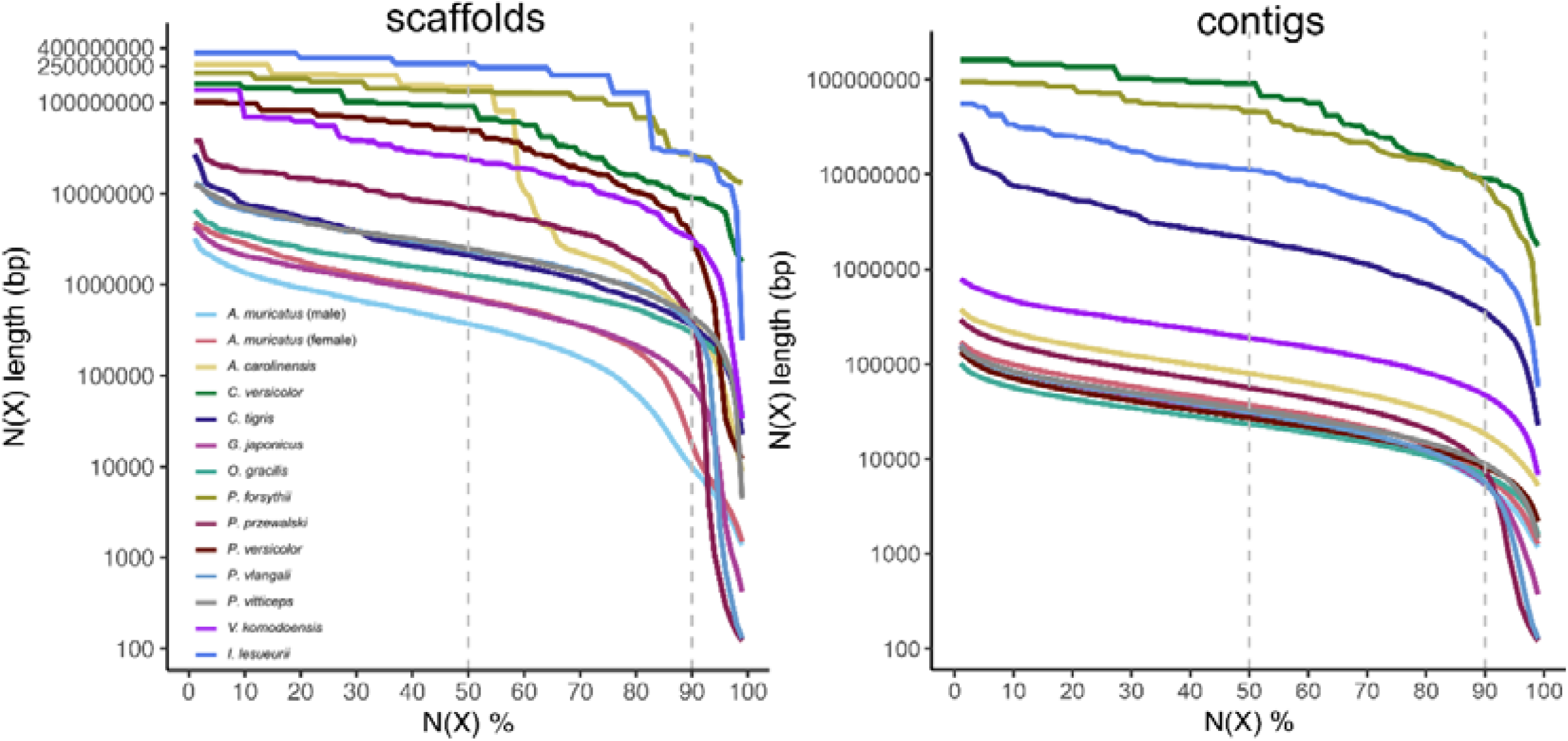
Comparison of the contiguity of two *A. muricatus* assemblies and 12 publicly available squamate assemblies. N(x)% graphs show the (A) contig and (B) scaffold lengths (*y-*axis), where x% (*x*-axis) of the genome assembly consist of scaffolds and contigs of at least that size. Dashed, grey lines denote N50 and N90 values. AmpMurF_3.0 and AmpMurM_3.0. denotes the female and male *A. muricatus* assembly, respectively

**Figure 3.**
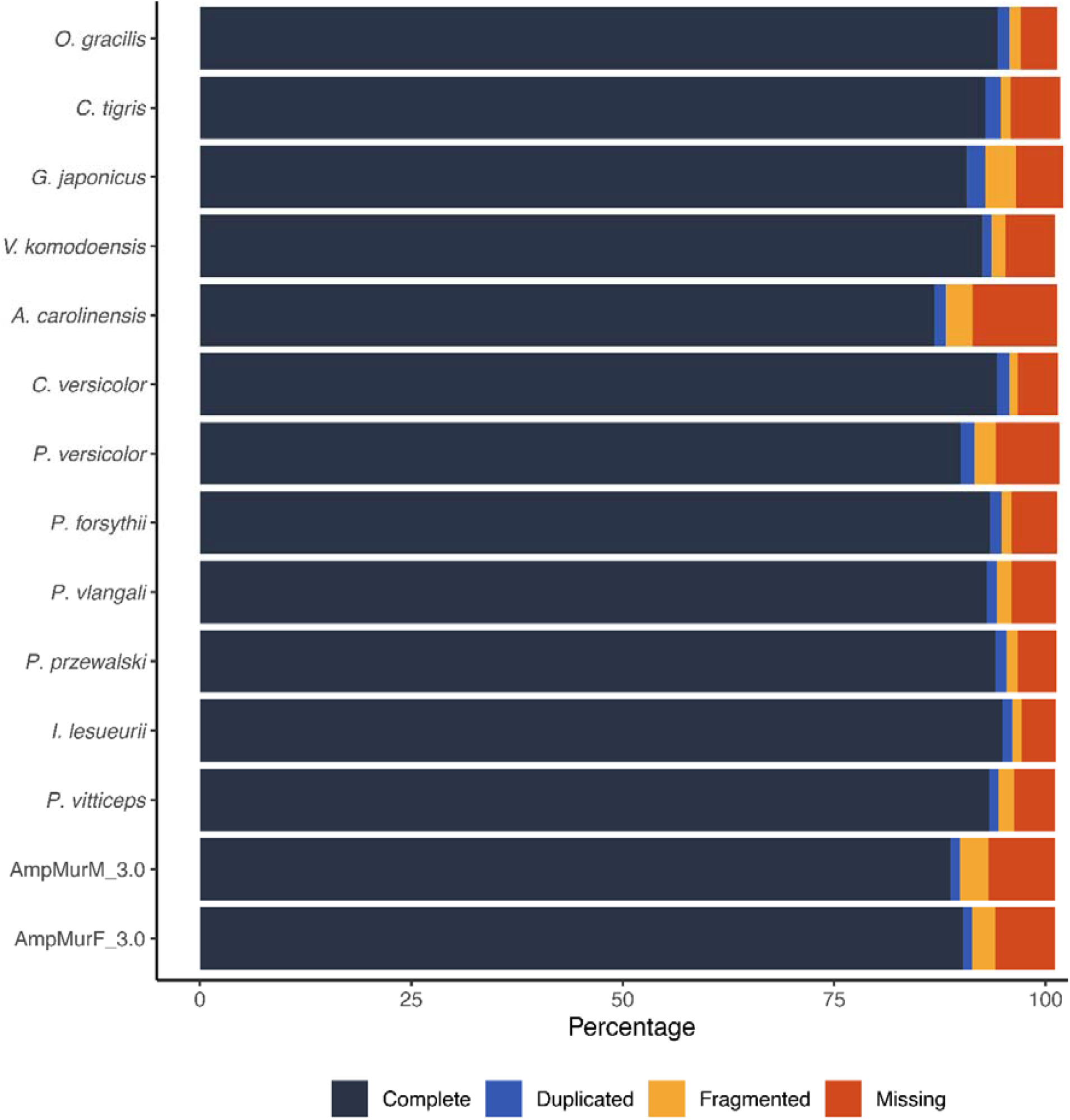
BUSCO assessment of genome assemblies from 13 squamate species. All assemblies were examined using the same version and library of BUSCO (v5.0.0 with the 7,480-gene sauropsida_odb10 dataset). AmpMurF_3.0 and AmpMurM_3.0. denotes the female and male *A. muricatus* assembly, respectively.

### Genome annotation

The *A. muricatus* assemblies are composed of ∼38% repeat elements and have a GC content of ∼42% (**Tables S6-S8**), similar to that of *P. vitticeps* (Georges et al. 2015) – with LINEs being the predominant subtype. Protein-coding genes were annotated by combining transcriptome evidence with homology-based (*A. carolinensis, V. komodoensis*, and *P. vitticeps*) and de novo gene prediction methods. Gene statistics (**Table S9**) (see (Georges et al. 2015)) and gene set BUSCO scores (**Table S10**) are comparable to other squamates. Using ab inito, transcriptome, and homology-based prediction methods, we functionally annotated 21,655 (95.12%) and 21,799 (94.70%) protein-coding genes in the female and male assembly (**Tables S11 and S12**) and recovered 89.2% and 88.2% of 7,480 sauropsid (i.e., non-avian reptiles and birds) benchmarking universal single-copy orthologs (BUSCOs), respectively.

### Phylogenetic relationships

To construct a time-calibrated species tree (**Figure 4**), we identified 3,130 single-copy orthologs from the female *A. muricatus* assembly and 12 other squamate species. In addition to *A. muricatus*, seven agamid species currently have genome assemblies: two Australian dragon lizards (*P. vitticeps* and *I. lesueurii*), four toad-headed agamas (genus *Phrynocephalus*), and the oriental garden lizard (*C. versicolor*) (Gao et al. 2019; Georges et al. 2015; Qi et al. 2023; Wang et al. 2023; Jin et al. 2022). Our analysis shows that the agamids shared an ancestor about 79.8 Mya [74.5–86.1 Mya 95% credibility interval (CI)]. We estimate that the three Australian dragon lizard species split from a common ancestor about 37.2 Mya (95% CI 30.6–42.2), while the lineages leading to *A. muricatus* and *P. vitticeps* diverged 24.6 Mya (95% CI 19.6–31.7). These observations agree with previous dating from a small set of genes and fossil data (Hugall et al. 2008; Oliver and Hugall 2017).

**Figure 4.**
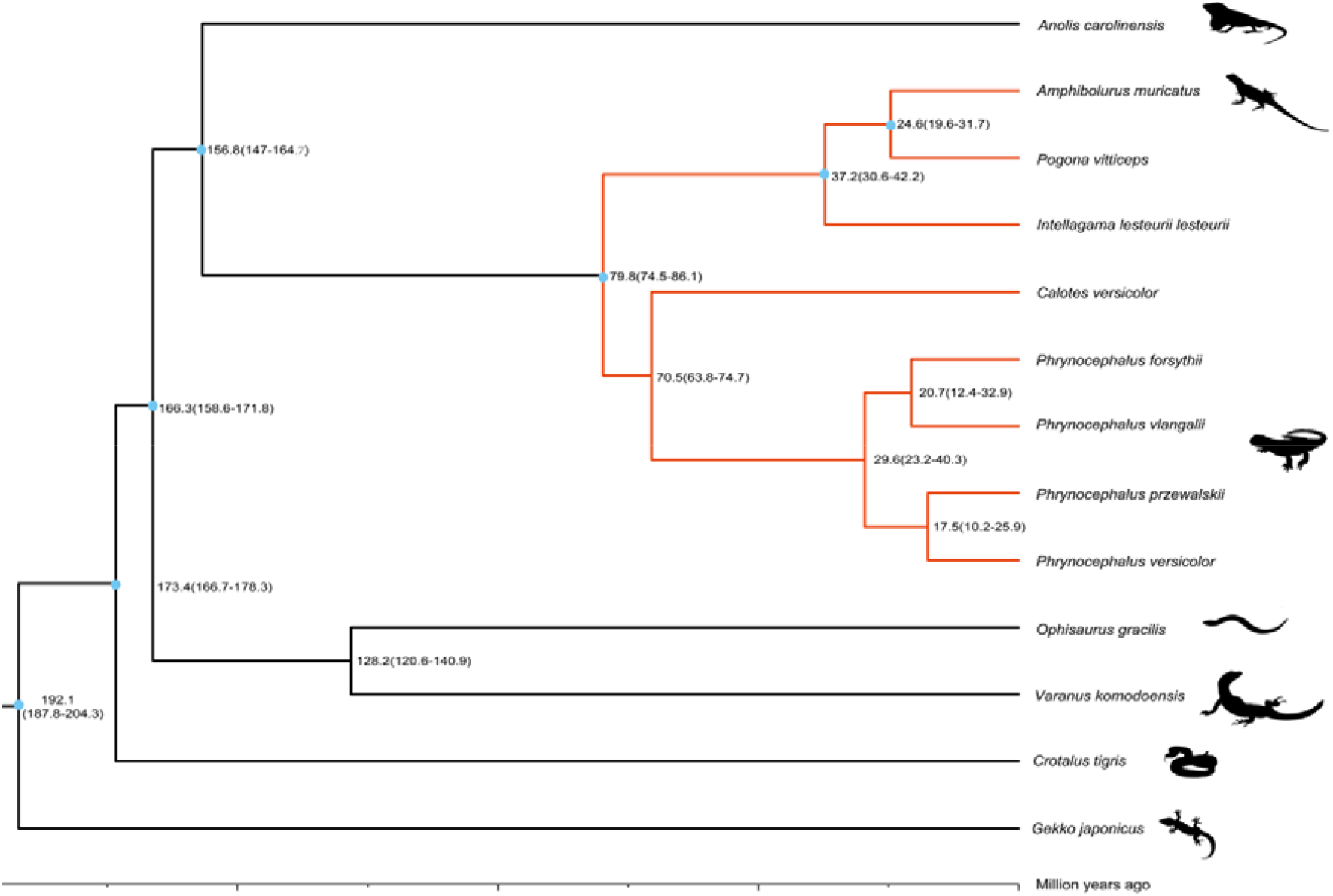
Inferred phylogeny of 13 squamate species based on whole-coding sequences of 3,130 1:1 orthologs. Numbers at nodes represent the estimated divergence time from present (million years ago; Mya) between lineages. Agamid (family Agamidae) lineages are indicated in red.

### Conclusions and perspectives

In this study, we generated the first annotated genome assemblies of *Amphibolurus muricatus*. Overall, the assemblies are similar in quality to a range of squamate genomes and will be immediately useful for the understanding of agamid lizard evolution, ecology, and conservation.

## Data availability

*A. muricatus* raw 10x Genomics genome and transcriptome sequencing reads have been deposited to the NCBI Short Read Database (BioProject ID: PRJNA767251). Raw stLFR genome sequencing reads have been deposited at the China National GeneBank Nucleotide Sequence Archive (CNSA: https://db.cngb.org/cnsa) under accession number CNP0004768. The male and female *A. muricatus* genome assemblies are available at Zenodo (Tian et al. 2023a). Gene annotation files and associated FASTA files for *A. muricatus* (assembly AmpMurF_3.0 and AmpMurM_3.0), *I. lesueurii, P. przewalskii*, and *P. vlangalii* are available at Zenodo (Tian et al. 2023b). *A. muricatus* transcriptome assemblies are available at Zenodo (Tian et al. 2021). Various scripts used for data processing and analyses are available on GitHub at https://github.com/sciseim/JackyDragon.

## Acknowledgements

We thank Dr Wendy Ruscoe and Jacqui Richardson for their assistance in generating the inbred line of *A. muricatus* and for animal husbandry.

## Ethics Approvals

All sampling and breeding experiments were conducted with approval of the Animal Ethics Committee of the University of Canberra and in accordance with their Standard Operating Procedures.

