## Supplemental Information for "Draft genomes of a male and female Australian jacky dragon (*Amphibolurus muricatus*)"

### **SUPPLEMENTARY FIGURES**

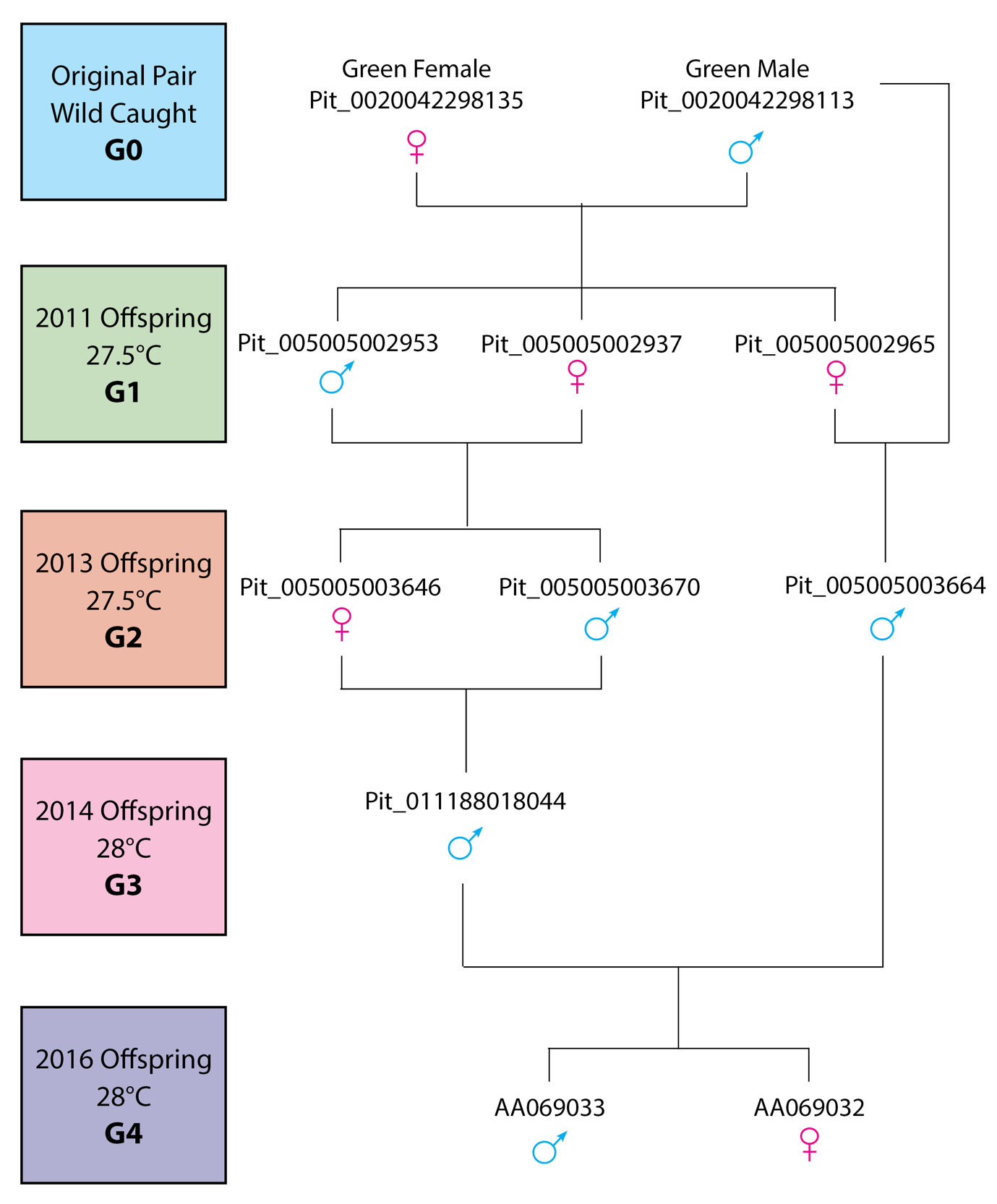

#### Figure S1 Pedigree for the inbred focal individuals used for genome sequencing (AA069032 and AA069033)

Part of the University of Canberra Wildlife Tissue Collection (NCBI Biocollections UC<Aus>).

### **SUPPLEMENTARY TABLES**

#### Table S1 *A. muricatus* transcriptome sequencing summary

Reads denotes million paired-end (PE) reads.

| Tissue | Sex | Sample code | Reads  (M) |
| --- | --- | --- | --- |
| Ovary | Female | AA069032 | 75.9 |
| Testis | Male | AA069033 | 72.2 |
| Brain | Male | AA069033 | 66.8 |
| Brain | Female | AA069032 | 66.3 |

#### Table S2 *A. muricatus* transcriptome assembly summary

Statistics are based on all transcript contigs. Total Trinity ‘genes’ refers to the number of transcript clusters generated by the assembly (isogroups), while Total Trinity Transcripts indicates the number of isoforms (isotigs).

|  | Tissue | | Total Trinity  'genes' | | Total Trinity 'transcripts' | | % GC | | N10 (bp) | | N50 (bp) | | Median contig length (bp) | | Average contig length (bp) | | Total  assembled bases (bp) | shortest contig  (bp) | | longest contig  (bp) |
| --- | --- | --- | --- | --- | --- | --- | --- | --- | --- | --- | --- | --- | --- | --- | --- | --- | --- | --- | --- | --- |
| Combined | | 497,966 | | 697,279 | | 43.91 | | 6,900 | | 1,889 | | 387 | | 876 | | 610,990,767 | | | 172 | 24,779 |
| Ovary | | 90,809 | | 136,091 | | 45.64 | | 7,300 | | 3,156 | | 481 | | 1,321 | | 179,750,369 | | | 184 | 22,312 |
| Testes | | 217,058 | | 329,033 | | 43.46 | | 4,959 | | 1,681 | | 414 | | 864 | | 284,237,931 | | | 179 | 15,438 |
| Brain | | 339,838 | | 443,832 | | 44.02 | | 6,978 | | 2,232 | | 378 | | 915 | | 406,213,309 | | | 180 | 24,870 |
| Female brain | | 182,133 | | 231,358 | | 44.62 | | 6,393 | | 2,311 | | 368 | | 925 | | 213,989,392 | | | 178 | 19,476 |
| Male brain | | 275,797 | | 357,624 | | 43.96 | | 7,090 | | 2,380 | | 378 | | 945 | | 338,062,929 | | | 183 | 27,615 |

#### Table S3 *A. muricatus* transcriptome assembly BUSCO statistics

Each transcriptome was assessed using BUSCO v5.0.0 and the 7,480-gene BUSCO data set sauropsida_odb10.

| Transcriptome | Complete  BUSCOs (C) | Fragmented BUSCOs (F) | Missing BUSCOs (M) |
| --- | --- | --- | --- |
| Combined | 6,952 (92.9%) | 155 | 373 |
| Ovary | 6,065 (81.1%) | 201 | 1,214 |
| Testis | 5,493 (73.5%) | 376 | 1,611 |
| Brain | 6,624 (88.6%) | 227 | 629 |
| Male brain | 6,448 (86.2%) | 240 | 792 |
| Female brain | 6,014 (80.4%) | 349 | 1,117 |

#### Table S4 *A. muricatus* genome assembly statistics.

Lengths in base pairs (bp). Note: assembly 1.0 denotes 10x Genomics Supernova; assembly 1.1, 10x Genomics Supernova + RNA read + Trinity scaffolding; assembly 2.0, 10x Genomics Supernova (assembly 1.0) + SLR-superscaffolder (with stLFR reads); assembly 3.0, stLFR Supernova. Unmasked assemblies were interrogated.

| **Assembly methods** | **Female (AmpMurF_1.0)** | **Female (AmpMurF_1.1)** | **Female (AmpMurF_2.0)** | **Female (AmpMurF_3.0)** | **Male (AmpMurM_1.0)** | **Male (AmpMurM_1.1)** | **Male (AmpMurM_2.0)** | **Male (AmpMurM_3.0)** |
| --- | --- | --- | --- | --- | --- | --- | --- | --- |
| Contig number | 154,897 | 124,200 | 154,961 | 145,095 | 180,498 | 151,787 | 180,576 | 95,472 |
| Contig length | 1,746,759,340 | 1,750,545,991 | 1,747,055,957 | 1,804,035,661 | 1,735,812,295 | 1,741,048,453 | 1,736,173,368 | 1,752,355,218 |
| Contig N50 (bp) | 25,056 | 37,220 | 25,053 | 67,166 | 21,019 | 28,761 | 21,019 | 59,294 |
| Contig max length | 209,568 | 348,284 | 209,568 | 645,479 | 196,238 | 288,200 | 196,238 | 773,372 |
| Scaffold number | 66,776 | 57,227 | 55,562 | 97,556 | 89,344 | 73,856 | 74,726 | 45,762 |
| Scaffold length | 1,840,499,790 | 1,841,491,868 | 1,871,715,150 | 1,868,109,324 | 1,831,120,515 | 1,833,283,242 | 1,854,210,455 | 1,804,786,947 |
| Scaffold N50 (bp) | 371,335 | 720,518 | 1,003,329 | 6,818,063 | 180,405 | 369,860 | 323,786 | 1,568,728 |
| Scaffold max length | 4,131,007 | 6,534,950 | 6,926,244 | 39,679,044 | 1,944,226 | 6,446,322 | 3,040,810 | 9,582,854 |
| Gaps (bp) | 93,740,450 | 90,945,877 | 124,659,193 | 64,073,663 | 95,308,220 | 92,234,789 | 118,037,087 | 52,431,729 |
| Gaps (%) | 5.09 | 4.94 | 6.66 | 3.43 | 5.20 | 5.03 | 6.37 | 2.91 |
| GC content (%) | 41.76 | 41.77 | 41.76 | 41.75 | 41.69 | 41.70 | 41.70 | 41.67 |

#### Table S5 BUSCO evaluation of squamate genome assemblies

For consistency, all genome assemblies were examined using the same version and library of BUSCO (5.0.0_cv1 with the 7,480-gene sauropsida _odb10 dataset). AmpMurF_3.0 and AmpMurM_3.0 denotes the female and male *Amphibolurus muricatus* assemblies, respectively. NA denotes not applicable. Assemblies with the suffix _HiC are available at DNA Zoo; CalVer_1.0 (CNP0003598), Phprz_v1.0 (CNP0000203), and Phvla_v1.0 (CNP0000203) are available at the China National GeneBank Database; the remainder are available via NCBI.

| Squamate family | Latin name | Common name | Assembly | Complete BUSCOs (C) | Complete and single-copy BUSCOs (S) | Complete and duplicated BUSCOs (D) | Fragmented BUSCOs (F) | Missing BUSCOs (M) |
| --- | --- | --- | --- | --- | --- | --- | --- | --- |
| Agamidae | *Amphibolurus muricatus* | Jacky dragon | AmpMurF_3.0 | 6,749 (90.2%) | 6,666 | 83 | 203 | 528 |
| Agamidae | *Amphibolurus muricatus* | Jacky dragon | AmpMurM_3.0 | 6,641 (88.8%) | 6,559 | 82 | 254 | 585 |
| Agamidae | *Pogona vitticeps* | Central bearded dragon | pvi1.1 | 6,982 (93.3%) | 6,899 | 83 | 138 | 360 |
| Agamidae | *Intellagama lesueurii* | Australian water dragon | EWD_hifiasm_HiC | 7,097 (94.9%) | 7,006 | 91 | 80 | 303 |
| Agamidae | *Phrynocephalus przewalskii* | Przewalski's toadhead agama | Phprz_v1.0 | 7,038 (94.1%) | 6,941 | 97 | 98 | 344 |
| Agamidae | *Phrynocephalus vlangalii* | Ching Hai toadhead agama | Phvla_v1.0 | 6,958 (93.0%) | 6,865 | 93 | 128 | 394 |
| Agamidae | *Phrynocephalus forsythii* | Moustache toadhead agama | ASM2928247v1 | 6,988 (93.4%) | 6,884 | 104 | 87 | 405 |
| Agamidae | *Phrynocephalus versicolor* | Variegated toadhead agama | PhrVer | 6,729 (89.9%) | 6,606 | 123 | 187 | 564 |
| Agamidae | *Calotes versicolor* | Oriental garden lizard | CalVer_1.0 | 7,048 (94.3%) | 6,939 | 109 | 78 | 354 |
| Agamidae | *Calotes versicolor* | Oriental garden lizard | ASM2071127v1 | 705 (9.5%) | 692 | 13 | 1,098 | 5,677 |
| Dactyloidae | *Anolis carolinensis* | Green anole | AnoCar2.0 | 6,495 (86.9%) | 6,394 | 101 | 238 | 747 |
| Varanidae | *Varanus komodoensis* | Komodo dragon | ASM479886v1 | 6,923 (92.6%) | 6,842 | 81 | 119 | 438 |
| Gekkonidae | *Gekko japonicus* | Schlegel's Japanese gecko | Gekko_japonicus_V1.1 | 6,786 (90.7%) | 6,627 | 159 | 276 | 418 |
| Viperidae | *Crotalus tigris* | Tiger rattlesnake | ASM1654583v1 | 6,948 (92.9%) | 6,815 | 133 | 89 | 443 |
| Anguidae | *Ophisaurus gracilis* | Anguidae lizard | NA | 7,058 (94.4%) | 6,957 | 101 | 100 | 322 |

#### Table S6 Repetitive sequence prediction of *A. muricatus* genomes

#### Female denotes assembly AmpMurF_3.0; male AmpMurM_3.0.

|  | **Female** | | **Male** | |
| --- | --- | --- | --- | --- |
| **Type** | **Repeat Size (bp)** | **% of genome** | **Repeat Size (bp)** | **% of genome** |
| TRF | 37,351,497 | 1.999 | 32,338,910 | 1.792 |
| RepeatMasker | 303,642,607 | 16.254 | 287,888,848 | 15.951 |
| Proteinmask | 219,444,832 | 11.747 | 210,064,729 | 11.639 |
| de novo | 700,617,740 | 37.504 | 669,397,681 | 37.09 |
| Total | 733,205,829 | 39.249 | 701,507,334 | 38.869 |

#### Table S7 Percentage of repetitive sequences in the female *A. muricatus* genome (assembly AmpMurF_3.0)

|  | **Repbase TEs** | | **Protein TEs** | | **de novo TEs** | | **Combined TEs** | |
| --- | --- | --- | --- | --- | --- | --- | --- | --- |
| **Type** | **Length(bp)** | **% of genome** | **Length(bp)** | **% of genome** | **Length(bp)** | **% of genome** | **Length(bp)** | **% of genome** |
| DNA | 50,755,532 | 2.717 | 1,906,049 | 0.102 | 15,844,709 | 0.848 | 62,525,206 | 3.347 |
| LINE | 210,520,993 | 11.269 | 185,692,304 | 9.94 | 551,394,796 | 29.516 | 595,778,878 | 31.892 |
| SINE | 21,913,866 | 1.173 | 0 | 0 | 5,531,160 | 0.296 | 26,748,749 | 1.432 |
| LTR | 33,518,243 | 1.794 | 31,931,219 | 1.709 | 154,264,576 | 8.258 | 170,999,054 | 9.154 |
| Other | 61,853 | 0.003 | 0 | 0 | 0 | 0 | 61,853 | 0.003 |
| Unknown | 0 | 0 | 0 | 0 | 10,943,014 | 0.586 | 10,943,014 | 0.586 |
| Total | 303,642,607 | 16.254 | 219,444,832 | 11.747 | 694,572,899 | 37.181 | 720,032,362 | 38.543 |

#### Table S8 Percentage of repetitive sequences in the male *A. muricatus* genome (assembly AmpMurM_3.0)

|  | **Repbase TEs** | | **Protein TEs** | | **de novo TEs** | | **Combined TEs** | |
| --- | --- | --- | --- | --- | --- | --- | --- | --- |
| **Type** | **Length(bp)** | **% of genome** | **Length(bp)** | **% of genome** | **Length(bp)** | **% of genome** | **Length(bp)** | **% of genome** |
| DNA | 46,637,125 | 2.584 | 2,842,746 | 0.158 | 16,067,062 | 0.89 | 59,181,370 | 3.279 |
| LINE | 200,767,375 | 11.124 | 177,004,166 | 9.807 | 544,149,840 | 30.15 | 576,905,171 | 31.965 |
| SINE | 21,052,333 | 1.166 | 0 | 0 | 5,817,210 | 0.322 | 25,898,781 | 1.435 |
| LTR | 30,426,547 | 1.686 | 30,280,042 | 1.678 | 134,392,379 | 7.446 | 149,836,069 | 8.302 |
| Other | 54,247 | 0.003 | 0 | 0 | 0 | 0 | 54,247 | 0.003 |
| Unknown | 0 | 0 | 0 | 0 | 5,271,406 | 0.292 | 5,271,406 | 0.292 |
| Total | 287,888,848 | 15.951 | 210,064,729 | 11.639 | 664,607,189 | 36.825 | 690,074,989 | 38.236 |

#### Table S9 Overview of *A. muricatus* gene sets

AmpMurF_3.0 and AmpMurM_3.0 denotes the female and male *Amphibolurus muricatus* assemblies, respectively.

|  | AmpMurF_3.0 | AmpMurM_3.0 |
| --- | --- | --- |
| No. protein-coding genes | 22,765 | 23,019 |
| Average gene length (bp) | 27,241.16 | 26,280.65 |
| Average CDS length (bp) | 1,428.76 | 1,404.82 |
| Average exon no. | 8.31 | 8.18 |
| Average exon length (bp) | 171.95 | 171.71 |
| Average intron length (bp) | 3,531.52 | 3,464.02 |

#### Table S10 BUSCO evaluation of squamate gene sets

Each gene set (CDS) was assessed using BUSCO v5.0.0 in transcriptome mode and the 7,480-gene BUSCO data set sauropsida_odb10. AmpMurF_3.0 and AmpMurM_3.0 denotes the female and male *Amphibolurus muricatus* assemblies, respectively.

| Squamate family | Latin name | Common name | Complete BUSCOs (C) | Fragmented BUSCOs (F) | Missing BUSCOs (M) |
| --- | --- | --- | --- | --- | --- |
| Agamidae | *Amphibolurus muricatus* | Jacky dragon (AmpMurF_3.0) | 6,674 (89.2%) | 219 | 587 |
| Agamidae | *Amphibolurus muricatus* | Jacky dragon (AmpMurM_3.0) | 6,601 (88.2%) | 268 | 611 |
| Agamidae | *Pogona vitticeps* | Central bearded dragon | 6,946 (92.9%) | 188 | 346 |
| Agamidae | *Intellagama lesueurii* | Australian water dragon | 5,930 (79.3%) | 239 | 1,311 |
| Agamidae | *Phrynocephalus przewalskii* | Przewalski's toadhead agama | 6,326 (84.5%) | 204 | 950 |
| Agamidae | *Phrynocephalus vlangalii* | Ching Hai toadhead agama | 6,218 (83.1%) | 249 | 1,013 |
| Agamidae | *Phrynocephalus forsythii* | Forsyth's toadhead agama | 6,257 (83.7%) | 336 | 887 |
| Agamidae | *Phrynocephalus versicolor* | Tuvan toad-headed agama | 6,841 (91.5%) | 229 | 410 |
| Agamidae | *Calotes versicolor* | Oriental garden lizard | 7,077 (94.6%) | 157 | 246 |
| Dactyloidae | *Anolis carolinensis* | Green anole | 7,009 (93.7%) | 88 | 383 |
| Varanidae | *Varanus komodoensis* | Komodo dragon | 7,222 (96.6%) | 52 | 206 |
| Gekkonidae | *Gekko japonicus* | Schlegel's Japanese gecko | 6,974 (93.3%) | 234 | 272 |
| Viperidae | *Crotalus tigris* | Tiger rattlesnake | 7,178 (96.0%) | 50 | 252 |
| Anguidae | *Ophisaurus gracilis* | Anguidae lizard | 5,034 (67.3%) | 800 | 1,646 |

#### Table S11 Functional annotation statistics of *A. muricatus* protein-coding genes

AmpMurF_3.0 and AmpMurM_3.0 denotes the female and male *Amphibolurus muricatus* assemblies, respectively. EVM denotes EVidenceModeler; KEGG, Kyoto Encyclopedia of Genes and Genomes; nr, NCBI non-redundant protein sequences.

|  | AmpMurF_3.0 |  | AmpMurM_3.0 |
| --- | --- | --- | --- |
| No. EMV gene models | 22,765 |  | 23,019 |
| nr | 20,096 (88.28%) |  | 20,206 (87.78%) |
| Swiss-Prot | 17,746 (77.95%) |  | 17,783 (77.25%) |
| KEGG | 17,609 (77.35%) |  | 17,638 (76.62%) |
| InterPro | 20,812 (91.42%) |  | 20,962 (91.06%) |
| Total annotated genes | 21,655 (95.12%) |  | 21,799 (94.70%) |

#### Table S12 List of functionally annotated *A. muricatus* protein-coding genes

AmpMurF_3.0 and AmpMurM_3.0 denotes the female and male Amphibolurus muricatus assemblies, respectively. NA denotes not applicable; EVM, EVidenceModeler; KEGG, Kyoto Encyclopedia of Genes and Genomes; nr, NCBI non-redundant protein sequences.

(in separate Excel file)
